# Sex-specific classification of drug-induced Torsade de Pointes susceptibility using cardiac simulations and machine learning

**DOI:** 10.1101/2020.10.26.352740

**Authors:** Alex Fogli Iseppe, Haibo Ni, Sicheng Zhu, Xianwei Zhang, Raffaele Coppini, Pei-Chi Yang, Uma Srivatsa, Colleen E. Clancy, Andrew G. Edwards, Stefano Morotti, Eleonora Grandi

**Author notes:** **Corresponding author**, Name: Eleonora Grandi, Address: Department of Pharmacology, University of California, Davis, CA, USA.

## Abstract

Torsade de Pointes (TdP), a rare but lethal ventricular arrhythmia, is a toxic side effect of many drugs. To assess TdP risk, safety regulatory guidelines require quantification of hERG channel block *in vitro* and QT interval prolongation *in vivo* for all new therapeutic compounds. Unfortunately, these have proven to be poor predictors of torsadogenic risk, and are likely to have prevented safe compounds from reaching clinical phases. While this has stimulated numerous efforts to define new paradigms for cardiac safety, none of the recently developed strategies accounts for patient conditions. In particular, despite being a well-established independent risk factor for TdP, female sex is vastly underrepresented in both basic research and clinical studies, and thus current TdP metrics are likely biased toward the male sex. Here, we apply statistical learning to synthetic data, generated by simulating drug effects on cardiac myocyte models capturing male and female electrophysiology, to develop new sex-specific classification frameworks for TdP risk. We show that (1) TdP classifiers require different features in females vs. males; (2) male-based classifiers perform more poorly when applied to female data; (3) female-based classifier performance is largely unaffected by acute effects of hormones (i.e., during various phases of the menstrual cycle). Notably, when predicting TdP risk of intermediate drugs on female simulated data, male-biased predictive models consistently underestimate TdP risk in women. Therefore, we conclude that pipelines for preclinical cardiotoxicity risk assessment should consider sex as a key variable to avoid potentially lifethreatening consequences for the female population.

## 1. Introduction

During drug development, promising therapeutic compounds are tested to evaluate their potential risk of inducing Torsade de Pointes (TdP), a specific form of polymorphic ventricular tachycardia that can precipitate ventricular fibrillation and cause sudden cardiac death.^1^ While TdP is a very rare adverse event, amounting to less than one case out of 100,000 exposures for some non-antiarrhythmic drugs, cardiac safety concerns have caused withdrawal from the market of several drugs, including antihistamines, antidepressants, chemotherapeutics, pain medications, that had been associated with TdP proclivity in patients.^2^ The most simple mechanistic explanation of torsadogenicity involves a reduction of the rapid delayed rectifier K^+^ current (I_Kr_), carried by the human Ether-à-go-go-Related Gene (hERG) channel, which importantly contributes to cardiac action potential (AP) repolarization.^3^ Pharmacological block of the hERG channel, which is a very promiscuous target interacting with cardiac and non-cardiac drugs, produces AP duration (APD) and QT interval prolongation, and leads to an increased susceptibility to pro-arrhythmic events. Based on this evidence, current safety regulatory guidelines require the measurement of hERG channel block *in vitro* and QT interval prolongation *in vivo* to estimate TdP risk.^4,5^ Since their adoption, these guidelines have successfully avoided that cardiotoxic drugs could endanger the welfare of people. However, it has also become apparent that these biomarkers are poor predictors of torsadogenic risk, and have in all probability prevented safe treatments from reaching the market.^6,7^

In response to this problem, recent efforts have led to several proposed new paradigms for the prediction of TdP. One notable example is the Comprehensive In Vitro Proarrhythmia Assay (CiPA) initiative, an international multi-group initiative by regulatory, industry, and academic partners including the US Food and Drug Administration. This paradigm relies upon the idea of combining *in vitro* studies to measure the drugs effects on each of the different types of ion channels and *in silico* models of cardiac myocyte electrophysiology to understand how these effects combine to influence cardiac function, thus creating a novel tool for TdP risk assessment of new drugs.^9^ Mathematical models of cardiac electrophysiology, in fact, make it possible to simulate extreme conditions with precision (e.g., dangerously supratherapeutic drug concentrations), and to obtain insights generally only accessible via animal experiments. Thus, computational approaches have become essential components of numerous strategies to predict torsadogenic risk.^10–12^ In addition, simulated measurements extracted from the biophysical model simulations can also be applied to machine learning (ML) pipelines, as demonstrated by the Sobie group,^13^ with the potential to bring out mechanistic insights buried in the data that could be otherwise ignored.

To our knowledge, however, no simulation-based approach has considered any risk factor for TdP in their predictive pipelines. An emblematic example is provided by known sex-differences: it is well established that women are more susceptible to TdP than men when treated with QT-prolonging drugs,^14,15^ suggesting that TdP risk classifiers could benefit from inclusion of biological sex as a predictive variable. However, female sex is highly underrepresented in both basic research^16^ and clinical studies^17^ involved in the drug development process, with important consequences for the identification of accurate TdP predictors. *In vitro* studies tend to use mostly male animals,^16^ raising concerns on the generalizability of findings to the whole population. This sex bias propagates onto the mathematical models of cardiac cells,^18^ which are parameterized based on male-dominated datasets. The issue of underestimating potential health risks for women is then aggravated by the fact that female sex is also underrepresented in clinical cohorts,^17^ making training of classifiers harder due to the lack of reliable ground truth data.

Here, we combined simulations of these mathematical models with machine learning to generate sexspecific TdP risk classifiers. We simulated the effects of 59 training drugs under different conditions using *in silico* models of human ventricular myocytes with sex-specific parameterizations.^19,20^ We fed the resulting high-dimensional datasets of simulated biomarkers to machine learning algorithms to generate male and female classifiers of torsadogenic risk. Finally, we evaluated the effects of using sex-specific models for risk prediction on a separated set of 36 drugs, which are deemed at intermediate risk of TdP. Our results show that TdP classifiers trained on sex-specific datasets identify distinct and not interchangeable sets of optimal features, suggesting potential different drivers of drug-induced arrhythmias, and that the use of sex-biased predictive models underestimates the torsadogenic risk of drugs with intermediate risk of TdP in females, which could potentially lead to life-threatening consequences for women.

## 2. Methods

A schematic diagram describing the complete workflow used in the generation of the ML classifiers for TdP risk is shown in **Fig 1**, and a detailed Methods section is provided in the Supplementary Materials. Source code and documentation are freely available at http://elegrandi.wixsite.com/grandilab/downloads and https://github.com/drgrandilab.

**Figure 1:**
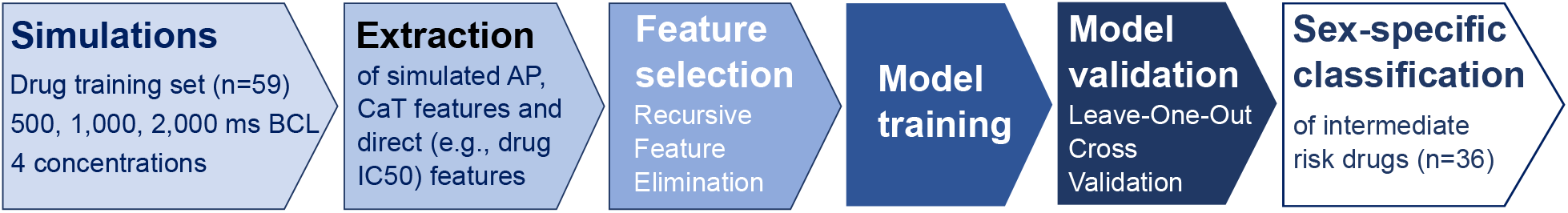
Workflow for the creation and testing of the sex-specific classifiers for torsadogenic risk.

### 2.1 Drug simulations

We updated the male and female human epicardial ventricular cardiomyocyte models developed by the Clancy lab,^20^ based on the O’Hara-Rudy model^21^ to recapitulate observed functional sex differences in Ca^2+^ handling.^22–24^ Drug effects were simulated with male and female models to build the complete set of biomarkers used to train the ML classifiers. The drug dataset used in this study (**Table S1**) was obtained by combining 83 compounds analyzed in the study by the Lancaster and Sobie with 12 CiPA compounds.^13,25^ Each drug was simulated using a pore block model based on the available IC_50_ values and Hill coefficients for various ion channels (a full list is available in **Table S1**). We included multiple concentrations, ranging from 1 to 4 times their effective free therapeutic plasma concentration (EFTPC), and various pacing rates, ranging from 0.5 to 2 Hz. We extracted 27 “simulated” biomarkers (**Table S2**) for each, and added “direct” measures, such as IC_50_ values for I_Kr_, I_Na_ (fast Na^+^ current) and I_CaL_ (L-type Ca^2+^ current).

### 2.2 Drug labels

Unfortunately, we are not aware of any clinical source of torsadogenic risk categorization that takes in account sex as an independent variable. In order to assign a unique binary label to each drug, we took advantage of the TdP risk classification available at www.crediblemeds.org,^26^ which reviews and analyzes adverse event reports for placing drugs in three broad categories: known, possible, and conditional risk of inducing TdP. Compounds characterized by known risk of inducing TdP in CredibleMeds were considered TdP^+^ in our analysis, while safe compounds (i.e., not included in any of the CredibleMeds categories) were considered TdP^-^. Given that sex is one of the potential risk factors for TdP, and thus determines the categorization of drugs in the possible or conditional risk categories, it would have been circular to assign these compounds to a specific class in our binary classification problem. Therefore, we did not use these drugs in training our classifiers. A similar selection was performed on the training set of drugs proposed by CiPA, which separates the compounds in high, intermediate, and low torsadogenic risk. Drugs classified differently by CredibleMeds and CiPA were excluded from the training dataset. Out of the 95 drugs in our initial dataset, 59 drugs met the requirements above and were used for creating our classifiers. The remaining 36 drugs, which are more likely to be associated with sex-specific effects, were later used to test our TdP classifiers and investigate the consequences of performing male-sex-biased vs. sex-specific predictions.

### 2.3 Machine Learning

In order to select the biomarkers contributing to the most accurate prediction of torsadogenic outcome, we applied a recursive feature elimination (RFE) algorithm, as detailed in the Supplementary Materials. For the sake of feature interpretability, we limited the ML modeling algorithms to logistic regression and support vector machine (SVM) with linear kernel. All the results shown here were obtained using SVM models, which outperformed logistic regression models on our datasets.

## 3. Results

### 3.1 Sex-specific biophysical models for action potential and Ca^2+^ transient

The Yang and Clancy model^20^ nicely recapitulates the prolonged AP observed in females vs. males.^27^ Conversely, we noted that the predicted larger Ca^2+^ loading and transient (CaT) amplitude in the female model does not match the sex differences measured in rodents.^24^ Our updated female model, with modifications in Ca^2+^-handling processes described in the Methods, captures these sex-specific differences (**Fig 2Aii**). Namely, the female CaT amplitude is modestly reduced (**Fig 2Biv**) and decays slightly more slowly (CaT duration at 50% of decay or CaD_50_, **Fig 2Bv**) than in male. At the same time, the diastolic [Ca^2+^] and sarcoplasmic reticulum (SR) content do not appreciably differ in male and female (**Fig 2Biii,Bvi**). These results are similar to Ca^2+^ measurements in myocytes from patients with cardiac hypertrophy (**Fig 2 *insets***), whereby no significant sex differences were detected in systolic or diastolic Ca^2+^ levels, decay rate, and SR Ca^2+^ content.^28–31^ Notably, AP properties are very modestly affected by the modifications we introduced in the model parameters, thus preserving the typical differences in repolarization between the two sexes in the original Yang and Clancy model (**Fig 2Ai,Bi,Bii**).^20^

**Figure 2:**
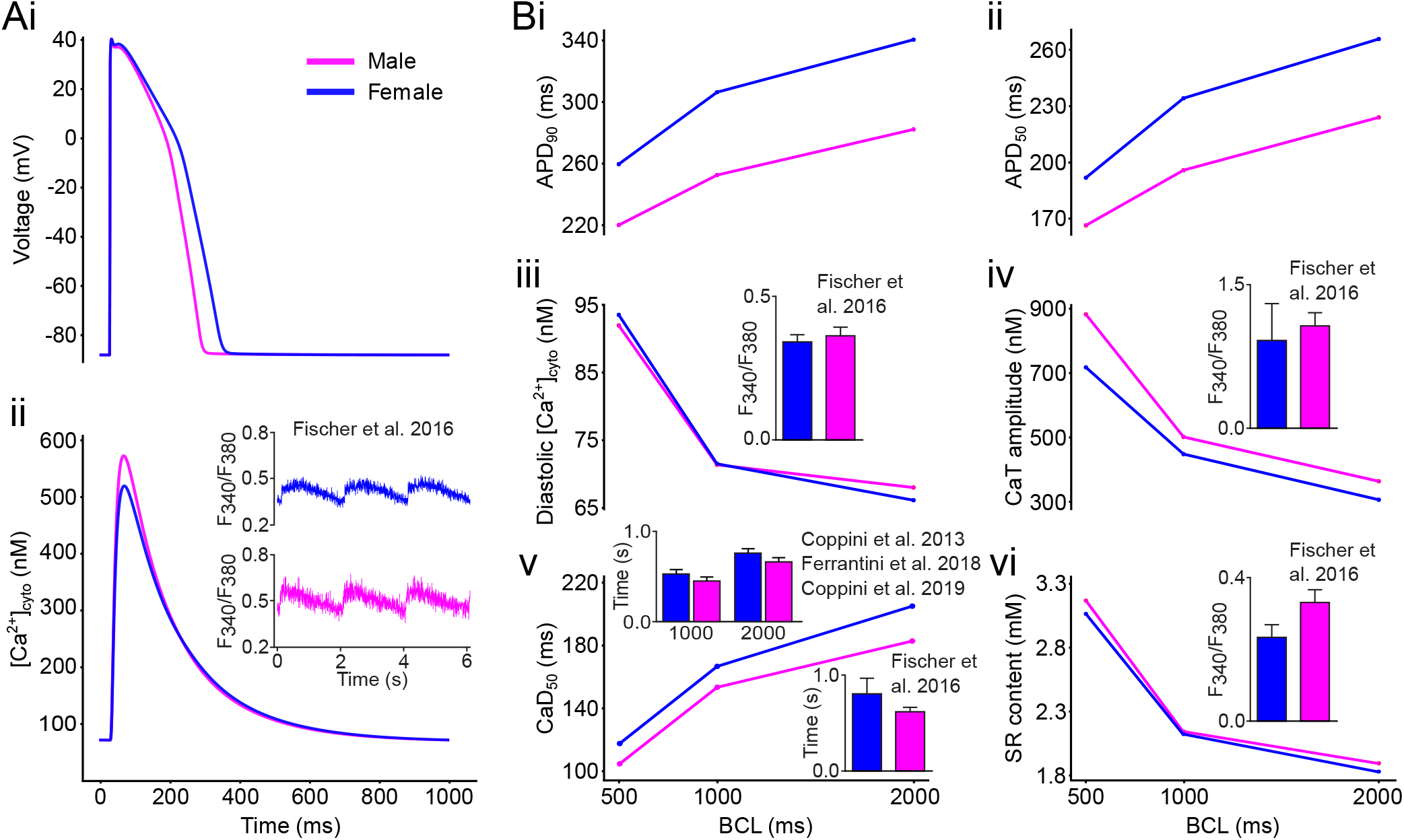
Sex-specific biophysical models of cardiac electrophysiology and Ca^2+^ handling. **A**: simulated AP (**i**) and CaT (**ii**) traces pacing at a BCL of 1,000 ms the baseline models of male (magenta) and female (blue) ventricular epicardial cardiomyocyte. **B**: Rate dependency of simulated biomarkers: measurements of voltage- (**i,ii**) and Ca^2+^-related (**iii-vi**) biomarkers as a function of the BCL tested. Insets show representative and summary data for CaT characteristics measured in myocytes from patients with cardiac hypertrophy paced at a BCL of 2,000 ms unless noted otherwise (reproduced with permission from Fischer et al. 2016,^28^ Coppini et al. 2013,^29^ Ferrantini et al. 2018,^30^ and Coppini et al. 2019^31^).

### 3.2 Sex-specific TdP risk classifiers

With the original male and updated female models, we simulated the effects of the 59 training drugs under different pacing and drug regimen conditions (see *Methods*). Representative simulated traces reflecting the ranges of variations induced by the drugs on the AP and CaT are shown in **Fig 3A** (male) and **B** (female). For each simulated condition, several biomarkers (**Table S2**) were extracted and, together with the IC_50_ values available for each drug, generated the final feature datasets used by the RFE algorithm. This process led to optimized male- and female-specific TdP classifiers.

**Figure 3:**
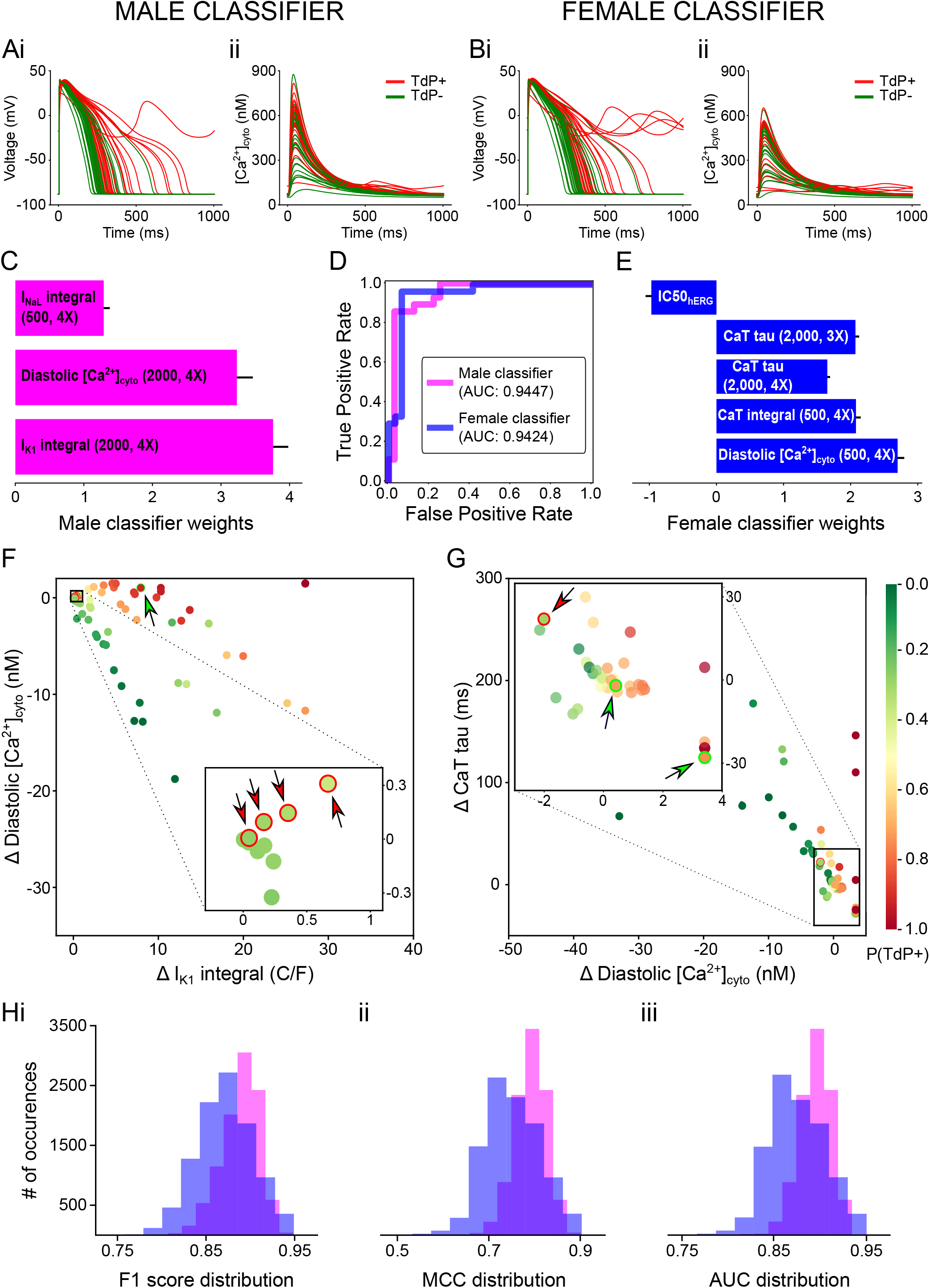
Sex-specific TdP risk classifiers. **A,B**: simulated effects of the 59 training drugs on the AP (i) and CaT (ii) with male (**A**) and female (**B**) biophysical myocyte models paced at a BCL of 1,000 ms. Traces belonging to drugs TdP^+^ and TdP^-^ are colored in red and green, respectively. **C,E**: best performing set of features selected through RFE and used to build the male (**C**) and female (**E**) TdP classifiers. Uncertainty of the feature weights is measured using LOO-CV (mean + SD). **D**: Receiver operating characteristic curve for male (magenta) and female (blue) TdP classifiers. Area under the curve (AUC) is 0.9447 and 0.9424 for male and female, respectively. **F,G**: scatterplot of the training drugs created using the two features with the largest weight for the male (**F**) and female (**G**) TdP classifiers. The estimated TdP risk probability for each drug is indicated by the color of the filling. The misclassified drugs are indicated by arrows, and the color of the arrow and the stroke specifies the right class (green for TdP^-^, red for TdP^+^). **H**: distributions of performance metrics for the male (magenta) and female (blue) TdP classifiers after injecting random normally-distributed (μ=0, σ=0.1) noise to the original data (10,000 repetitions).

The final sets of features and relative weights selected and used by the best performing male and female TdP risk classifiers are illustrated in **Fig 3C** and **3E**, respectively. TdP risk predictions for the male dataset are based on the current integrals of the inward rectifier I_K1_ and late Na^+^ current I_NaL_, and the di-astolic [Ca^2+^]. With LOO-CV, this ML model can correctly classify 54 out of 59 drugs, where-by 4 proarrhythmic drugs are predicted safe (4 false negatives, FNs – Amiodarone 1, Amiodarone 2, Cilostazol, Donepezil, where Amiodarone is simulated using different available IC_50_ values as shown in **Table S1**) and 1 safe drug is predicted harmful (1 false positive, FP – Prenylamine). The best performing female classifier is built on a set of 5 features. Four of the features are measurements extracted from the CaT (decay times, integral, and diastolic [Ca^2+^]), while the fifth one is the value of IC_50_ for the hERG channel. The predictive model optimized on the female dataset misclassifies 3 drugs: 1 FN (Procainamide) and 2 FPs (Ajmaline, Prenylamine). Despite the different set of features involved, both classifiers clearly demonstrate good ability in distinguishing torsadogenic from safe drugs. To evaluate model performances with commonly used metrics, we calculated F1 scores of 0.9148 and 0.9492, and MCCs of 0.8336 and 0.8987 for male and female TdP risk classifier, respectively (**Table 1**). Robust performances were confirmed by AUC values of 0.9447 and 0.9424 (**Fig 3D**) values for male and female classifier, respectively.

**Table 1:** Summary of prediction performances for various combinations of TdP classifiers and feature datasets.

|  | **F1 score** | **MCC** | **AUC** | **FN** | **FP** | **Misclassified drugs** |
| --- | --- | --- | --- | --- | --- | --- |
| **M on M** | 0.9148 | 0.8336 | 0.9447 | 4 | 1 | Amiodarone 1, Amiodarone 2, Cilostazol, Donepezil, Prenylamine |
| **F on F** | 0.9492 | 0.8987 | 0.9424 | 1 | 2 | Ajmaline, Prenylamine, Procainamide |
| **M on F** | 0.7693 | 0.5945 | 0.9159 | 12 | 1 | Amiodarone 1, Amiodarone 2, Bepridil 2, Bepridil CiPA, Cilostazol, Donepezil, Flecainide, Halofantrine, Prenylamine, Procainamide, Quinidine 1, Sotalol, Thioridazine 1 |
| **M retrained on F** | 0.8980 | 0.7972 | 0.9217 | 4 | 2 | Ajmaline, Amiodarone 1, Amiodarone 2, Cilostazol, Donepezil, Prenylamine |
| **F on EF** | 0.9322 | 0.8665 | 0.9516 | 1 | 3 | Ajmaline, Prenylamine, Procainamide, Telbivudine |
| **F on LF** | 0.9492 | 0.8987 | 0.9516 | 1 | 2 | Ajmaline, Prenylamine, Procainamide |
| **F on LU** | 0.9492 | 0.8987 | 0.9505 | 1 | 2 | Ajmaline, Prenylamine, Procainamide |

To visualize the TdP risk predictions of the classifiers, we created 2D scatterplots of the training compounds using the two most influential features used by each model (**Fig 3F** for male, **Fig 3G** for female). In this representation, the farther the features from the drug-free condition at coordinates (0,0), i.e., the larger the drug effect on the features, the more extreme are the predicted probabilities (darker colors). Interestingly, most of the misclassified drugs are located in the features space around the drug-free condition (insets of **Fig 3F** and **3G**), and are associated with probabilities close to the classification threshold of 0.5. To evaluate the robustness of our classifiers, we added random normally distributed noise (μ=0, σ=0.1) to the features and evaluated their performances by repeating this procedure 10,000 times (**Fig 3H**). The noise injection tends to increase the misclassification rate of the classifiers, whereby the drugs that are more frequently misclassified are those located in close proximity to the decision boundary of the SVM (r = -0.7653 and -0.8808 for male and female, respectively). However, despite being negatively affected, the performances of the sex-specific classifiers remain satisfactory even in presence of confounding noise in the data.

### 3.3 Testing the male classifier on female data

To verify our contention that the creation of sex-specific classifiers is indeed critical to obtain accurate predictions, we evaluated the performance of the male classifier in predicting TdP risk of the training drugs applied to the female simulated features. When applied to female data, the predictive accuracy of the male classifier dropped, producing 1 FP and 12 FNs (**Fig 4A**). One possible explanation for this poor performance could be that the male classifier was trained on a different dataset. To test this idea, we retrained the male classifier using the data generated from the simulations with the female biophysical model (weights are compared in the insert of **Fig 4B**) and re-evaluated its performance. The retrained male ML model still produced more misclassified drugs compared to our female classifier (**Fig 4A**). This result demonstrates that the set of features used by the female classifier has superior predictive ability for drug-induced arrhythmias in females, and confirms the need for considering sex when estimating drug-induced torsadogenic risk in preclinical compounds (and in patients).

**Figure 4:**
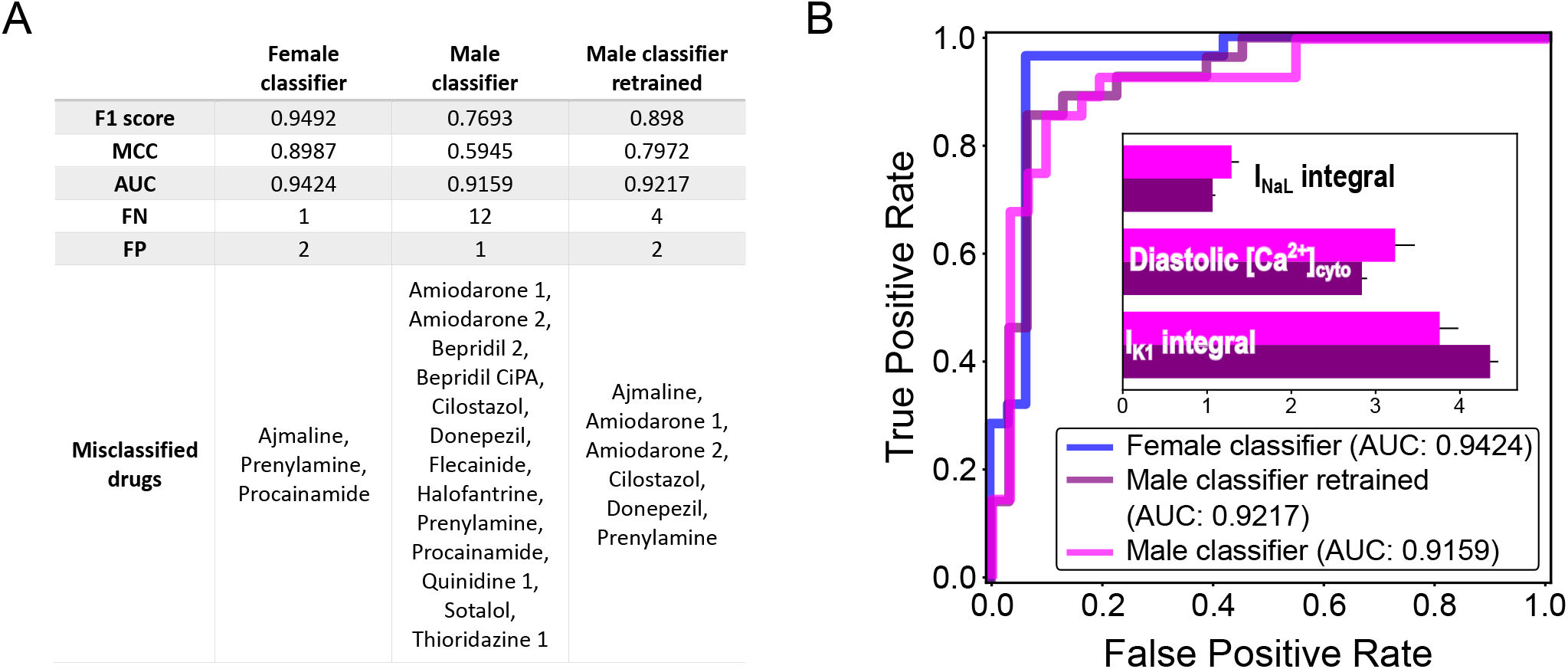
Predictions of TdP risk on female data with male-specific features. **A**: TdP classifier performances are evaluated on the dataset created with the female biophysical model. The results obtained with the female classifier (left column) are compared with the ones of the original male classifier (center column) and of a male classifier retrained on the female dataset (right column). **B**: Receiver operating characteristic curve for female (blue), original male (blue), and retrained male (purple) TdP classifiers applied on female data; area under the curve (AUC) is 0.9424, 0.9159, and 0.9217, respectively. In the inset, bar plot comparing the weights of the original (magenta) and retrained (purple) male classifiers. Uncertainty of the feature weights is measured using LOO-CV (mean + SD).

### 3.4 Sex-specific prediction of drugs with intermediate risk of TdP

From the initial list of drugs in our possession, 36 drugs had not been included in the training phase for belonging to the so-called intermediate torsadogenic risk category. In these drugs, TdP risk is associated with the presence of one or more risk factors. Indeed, when simulating these compounds with the sexspecific cardiomyocyte models, the observed changes on the AP (**Fig 5Ai,Bi**) and CaT (**Fig 5Aii,Bii**) are milder compared to the drugs with higher risk (**Fig 3A,B**). In order to explore how sex could affect the estimated torsadogenic risk of the drugs in the intermediate category, we applied the sex-specific TdP classifiers to the simulated male and female features and compared the predicted TdP probabilities (**Fig 5Ci**). In general, intermediate TdP risk drugs tend to have more dangerous outcomes in women, whereby a larger number of compounds is predicted to have higher probability of TdP in women (**Fig Cii**). Notably, if the male classifier is used on female data (right columns, **Fig 5Ci**), the predicted risk of the compounds is consistently underestimated, confirming the results obtained with the training dataset.

**Figure 5:**
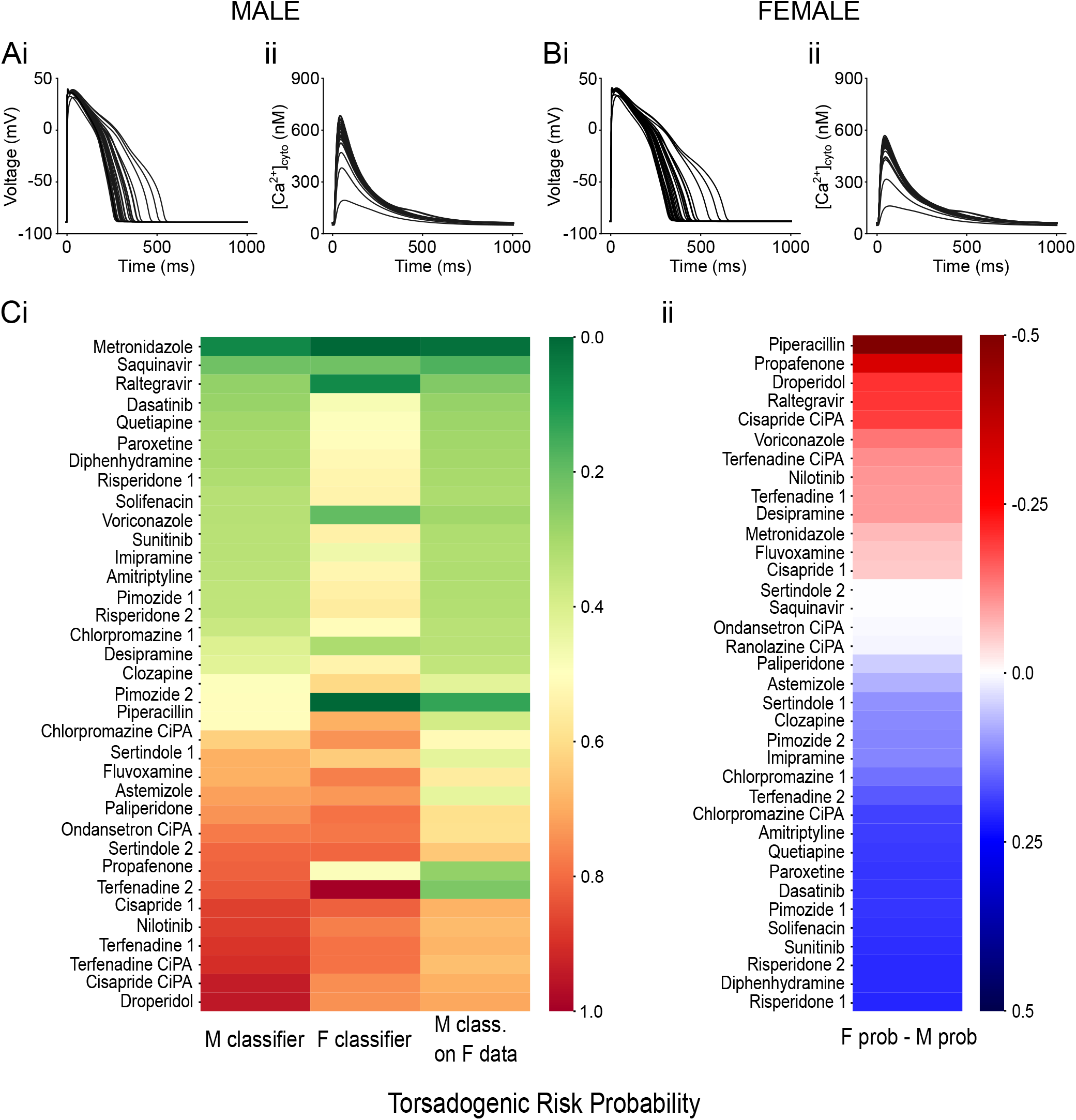
Sex-specific TdP predictions for intermediate-risk drugs. **A,B**: simulated effects of the 36 intermediate drugs on the AP (i) and CaT (ii) on the male (**A**) and female (**B**) biophysical models paced at a BCL of 1,000 ms. **C**: heatmaps of the predicted torsadogenic risk probability estimated by the sex specific TdP classifiers. (**i**) Left and center columns show the predictions obtained in male and female, respectively. The torsadogenic risk probabilities predicted by the male TdP classifier on the female dataset are visualized in the right column. Drugs are sorted by the risk probability predicted by the male TdP classifier. (**ii**) Difference between torsadogenic risk probabilities predicted by female and male TdP classifiers.

### 3.5 Effect of hormones on TdP risk prediction

It is well-established in the literature that circulating levels of hormones affect cardiac electrophysiology.^32^ To test how the performance of the female TdP classifier is influenced by hormones, we simulated the drug effects during the different phases of the menstrual cycle. Notably, the performance of the female classifier is almost unaltered by the hormonal effects (**Fig 6A**), with some modest changes in prediction in the late follicular phase. This is explained by hormones having minimal consequences on the CaT, which strongly influences the female predictions, while prolonging the AP duration considerably (**Fig 6B**). The consistency of the female classifier predictions in response to hormonal perturbations is also reflected in the probabilities forecasted for the drugs at intermediate TdP risk (**Fig 6C**). Similar results were obtained for the male classifier simulating the effects of testosterone (not shown). Taken together, these results show that this torsadogenic classifier is robust to acute changes in the levels of the sex hormones.

**Figure 6:**
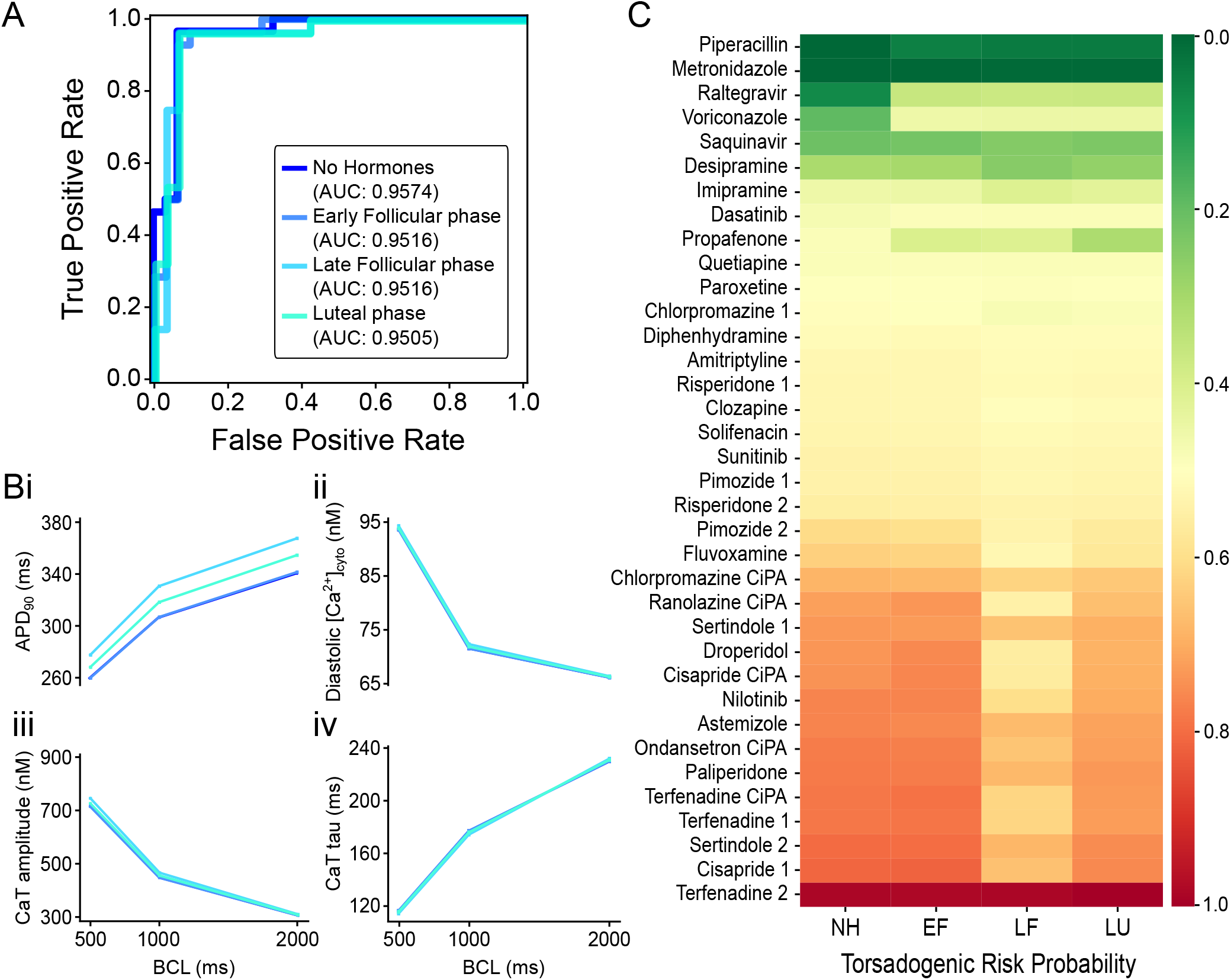
Effects of sex hormone fluctuations on female TdP classifier predictions. **A**: Receiver operating characteristic curve for female TdP classifier applied on training drugs dataset in absence of hormonal effects (navy blue) or simulating early follicular (EF, azure blue), late follicular (LF, sky blue), and luteal (LU, turquoise) phases. Area under the curve (AUC) is 0.9574, 0.9516, 0.9516, and 0.9505, respectively. **B**: Rate dependency of biomarkers simulated during different menstrual phases: measurements of voltage- (**i**) and Ca^2+^-related **(ii-iv**) biomarkers as a function of the BCL tested. **C**: heatmap of the predicted torsadogenic risk probabilities of intermediate TdP risk drugs simulated during different menstrual phases.

## 4. Discussion

Despite its important role in susceptibility to Torsade de Pointes,^14,15^ sex is rarely considered when developing predictive frameworks for torsadogenic risk. In the current study, we updated a published model of the human ventricular epicardial myocyte with sex-specific parameterizations^20^ to include experimentally observed differences in Ca^2+^ handling,^22–24^ and used it to simulate the effects of drugs belonging to different TdP risk categories. We then fed the simulated data to machine learning algorithms and generated sex-specific TdP risk classifiers. We showed that (1) classifiers trained on data reflecting male and female electrophysiological properties are built on distinct and not interchangeable sets of features; (2) male classifiers underestimate the torsadogenic risk in females; (3) female classifier predictions are robust to changes in sex hormone fluctuations during the menstrual cycle. Taken together, these results confirm the need for including sex when estimating the torsadogenic risk of a drug and provide a new tool to aid in this investigation.

### 4.1 Sex-specific model of ventricular electrophysiology and Ca^2+^ handling

The female Yang and Clancy^20^ model was parameterized using differences in expression levels of ion channels and transporters measured in cardiac tissue explanted from female vs. male patients, and included the effects of sex hormones on three ionic currents (I_Kr_, I_Ks_ and I_CaL_).^33^ While the resulting parameterizations accurately recapitulate the well documented differences in AP (and QT) properties in the basic research and clinical literature,^27,34^ less is known about sex differences in Ca^2+^ handling in humans. Experimental studies in rodents revealed smaller CaT amplitudes in female animals, associated with a decreased excitation-contraction coupling gain.^24^ Interestingly, L-type Ca^2+^ current, diastolic [Ca^2+^] and SR Ca^2+^ content are not significantly different between sexes.^24^ The simulated Ca^2+^ levels of the original female model were unusually high, particularly at faster frequencies and compared to the simulated male Ca^2+^ levels. Therefore, we modified some parameters affecting the Ca^2+^-handling processes in the female model (see Detailed Methods in the Supplementary Materials) and obtained CaTs in line with the differences reported in literature. Intriguingly, the outputs of our updated model more closely resemble the sex-specific Ca^2+^ recordings and related biomarkers acquired from isolated ventricular myocytes of hypertrophic patients (**Fig 2**).^28–31^ We did not detect sex differences in a small subset of control patients from previous publications (3 female vs 5 males, not shown).^29^ Information on the sexspecific differences in Ca^2+^ handling from a bigger sample of non-diseased patients would be highly desirable, but we are not aware of any published data with larger cohorts.

### 4.2 Sex-specific TdP biomarkers

Our work is preceded by a number of important studies that utilized *in silico* approaches to identify biomarkers predictive for TdP.^9–13,35^ Here we propose for the first time the adoption of sex-specific torsadogenic risk predictors. Through an RFE algorithm, we obtained male and female classifiers built on distinct minimal sets of best performing features. Importantly, the RFE iterative process does not require any manual intervention, removing opportunity for user bias. From the analysis of predictive biomarkers (**Fig 3C**), we found that the male classifier resembles the SVM classifier developed by Lancaster and Sobie.^13^ In both cases, TdP classification is based on changes in (1) diastolic [Ca^2+^] and (2) one or more metrics related to AP prolongation. Diastolic [Ca^2+^] is an index of cell Ca^2+^ loading. In fact, in these models it is highly correlated with SR Ca^2+^ content and CaT amplitude (r = 0.9803 and 0.8476, respectively). The integral of I_K1_ used by our male classifier is correlated with APD_90_/AP triangulation (r = 0.8872 and 0.8946 respectively). Accordingly, our **Fig 3F** is strikingly similar to Fig 3b in Lancaster and Sobie, 2016.^13^ The female classifier, on the contrary, selected hERG IC_50_ and several features related to Ca^2+^-handling processes. Diastolic [Ca^2+^], the integral of the CaT, and CaT decay time are all positively associated with torsadogenic risk (**Fig 3E**). Altered Ca^2+^ homeostasis has been shown to be an important determinant of TdP susceptibility. Increased whole-cell Ca^2+^ load is well known to be capable of destabilizing repolarization through enhanced forward mode I_NCX_.^36^ This effect could be exacerbated in women, for whom NCX expression is higher in physiological conditions.^37^ While a detailed mechanistic interpretation of the different CaT-related features for TdP remains unclear, their importance in this work clearly suggests that further investigation is warranted.

### 4.3 Ground truth classification

To solve the critical lack of ground truth regarding clinical sex-specific TdP risk classification, we operated under the assumption that drugs considered safe (i.e., missing in the CredibleMeds database,^26^ which contains a continuously updated database of therapeutic compounds categorized for torsadogenic risk) are so in both sexes. Conversely, drugs clearly associated with TdP (i.e., “Known Risk of TdP” category), are likely to be so in both sexes. On the contrary, drugs with intermediate risk of causing TdP (i.e., “Possible” or “Conditional Risk of TdP” categories) are more likely to change risk categorization depending on sex. Those drugs were therefore not used to build our classifiers.

Based on this assumed ground truth, our sex-specific TdP classifiers have shown remarkable predictive power (**Fig 3D,H, Table 1**). Additionally, looking further at the misclassified drugs, we note that both male and female ML models erroneously classified prenylamine. This compound, missing in the CredibleMeds^26^ and CiPA^8^ databases, is labeled as safe in our pipeline. However, prenylamine has been reported to cause TdP in patients,^38^ hence it is labeled as torsadogenic by other categorizations.^39^ This suggests that the classifiers in fact correctly detected the torsadogenicity of this compound despite an incorrect label. A similar conclusion can be reached exploring the clinical literature for ajmaline.^40^

### 4.4 Predictions on intermediate drugs

Our classification results with the intermediate risk drugs identified multiple therapeutic compounds that are predicted safe in men and torsadogenic in women (**Fig 5C**), which is in general agreement with the increased susceptibility to TdP in females. As drug-induced TdP is a rare event, prospective studies to evaluate the TdP risk factors are difficult to design and would require very large patient cohorts. We analyzed retrospective reviews of case reports, and notably, found a nice correspondence in the predictions for some specific compounds with observations published in the clinical literature. Indeed, female sex is a risk factor for documented TdP episodes associated with the use of Risperidone,^41^ Sunitinib,^42^ Paroxetine,^43^ and Quetiapine,^44^ which are among the intermediate-risk therapeutic compounds with the largest selectivity for women based on our estimated torsadogenic risk.

It is also important to understand sex-differences in TdP outcome in high-risk drugs, which also demonstrate female sex-prevalence, as demonstrated for quinidine, amiodarone, sotalol, disopyramide, bepridil, prenylamine (but not procainamide).^14^

### 4.5 Limitations

It is important to recognize that even at comparable drug dosages, drug exposure may vary between women and men owing to differences in absorption, distribution, metabolism and excretion that could explain higher TdP risk in women. The increased risk is not fully explained by sex difference in drug plasma levels,^45^ though cellular concentrations of a same systemic drug dose can vary across individuals and between sexes. Thus, future studies should account for both (population) pharmacokinetic and pharmacodynamic drug interactions. Experimental differences in pharmacokinetics observed between men and women have frequently been attributed to bodyweight differences and thus might be addressed by appropriate adjustment of dosage by body weight. In fact, in a cohort of more than 200 patients a statistically significant prevalence of dofetilide dose reduction or discontinuation was found in female vs. male patients mostly due to QTc prolongation, although no TdP cases were reported.^46^ On the other hand, the occurrence of TdP was not associated to any critical serum drug level of quinidine.^47^

### 4.6 Summary and future directions

Our results indicate the need for considering sex when developing and applying torsadogenic classifiers to obtain more accurate predictions and provide safer therapeutic treatments to patients. Of note, the current structure and size of the human thorough QT/QTc study for evaluation of proarrhythmic potential are prohibitive of sex-specific safety screening, as guidelines stipulate that only if a sign of harm is detected in a small (8-10 subjects) Phase I safety trial, then special populations (including women) should be assessed. We argue that the requirement for suitably powered sex-specific risk needs to be encoded in the early phase thorough QT/QTc study, and might be an attainable goal.

Clinical risk assessment and trials suggest that, besides sex, other patient conditions, i.e., age, disease, electrolyte imbalance, interaction with other drugs should all be taken into account in evaluating TdP risk.^48^ Data gathered from experiments in human induced pluripotent stem cells-derived cardiomyocytes obtained from male and female cell lines have been recently proven useful for investigating the sex-differences in torsadogenicity,^49^ suggesting that these patient-derived cells could be used to guide new models and paradigms for safety pharmacology accounting for patient conditions.

## 5. Study Highlights

### What is the current knowledge on the topic?

While female sex is a well-known risk factor for Torsade de Pointes (TdP), it is generally ignored by current safety regulatory guidelines and most of the proarrhythmia risk prediction models.

### What question did this study address?

We sought to develop and evaluate sex-specific TdP risk classifiers using mechanistic models of male and female cardiac myocytes electrophysiology.

### What does this study add to our knowledge?

We found that sex-specific TdP risk classification relies on distinct and not interchangeable sets of biomarkers and is robust to fluctuations in sex hormones levels.

### How might this change clinical pharmacology or translational science?

Accounting for sex-specific TdP risk is critical for pro-arrhythmic risk evaluation during drug development and can contribute to safer therapeutics.

## 6. Acknowledgements

This work was supported by the National Institutes of Health (NIH) Stimulating Peripheral Activity to Relieve Conditions grant 1OT2OD026580 (EG and CC); the National Heart, Lung, and Blood Institute grants R01HL131517, R01HL141214, P01HL141084 (EG), and R00HL138160 (SM); the American Heart Association Postdoctoral Fellowship 20POST35120462 (HN), Predoctoral Fellowship 20PRE35120465 (XZ), and grant 15SDG24910015 (EG); the UC Davis School of Medicine Dean’s Fellow award (EG) and Academic Senate grant (EG and US); the Health and Environmental Sciences Institute grant U01 FD006676-01 (AE).

## 7. Author Contributions

A.F.I. and E.G. wrote the manuscript; A.F.I., A.G.E., S.M. and E.G. designed the research; A.F.I. performed the research; A.F.I., H.N., S.Z., X.Z., R.C., U.S., A.G.E., S.M. and E.G. analyzed the data; A.F.I., H.N., P.Y., C.E.C. and E.G. contributed to new analytical tools.

## References

1. Dessertenne, F. La tachycardie ventriculaire a deux foyers opposes variables. Arch. Mal. Coeur Vaiss. 59, 263–272 (1966).

2. Haverkamp, W. et al. The potential for QT prolongation and proarrhythmia by non-antiarrhythmic drugs: clinical and regulatory implications. Report on a Policy Conference of the European Society of Cardiology. Eur. Heart J. 21, 1216–1231 (2000).

3. Sanguinetti, M. C., Jiang, C., Curran, M. E. & Keating, M. T. A mechanistic link between an inherited and an acquird cardiac arrthytmia: HERG encodes the IKr potassium channel. Cell 81, 299–307 (1995).

4. Food and Drug Administration International Conference on Harmonisation; Guidance on S7B Nonclinical Evaluation of the Potential for Delayed Ventricular Repolarization (QT Interval Prolongation) by Human Pharmaceuticals; Availability. Fed. Regist. 70, 61133–4 (2005).

5. Food and Drug Administration International Conference on Harmonisation; guidance on E14 Clinical Evaluation of QT/QTc Interval Prolongation and Proarrhythmic Potential for Non-Antiarrhythmic Drugs; availability. Notice. Fed. Regist. 70, 61134–5 (2005).

6. Sager, P. T. Key clinical considerations for demonstrating the utility of preclinical models to predict clinical drug-induced torsades de pointes. Br. J. Pharmacol. 154, 1544–1549 (2008).

7. Gintant, G. An evaluation of hERG current assay performance: Translating preclinical safety studies to clinical QT prolongation. Pharmacol. Ther. 129, 109–119 (2011).

8. Sager, P. T., Gintant, G., Turner, J. R., Pettit, S. & Stockbridge, N. Rechanneling the cardiac proarrhythmia safety paradigm: A meeting report from the Cardiac Safety Research Consortium. Am. Heart J. 167, 292–300 (2014).

9. Li, Z. et al. Assessment of an In Silico Mechanistic Model for Proarrhythmia Risk Prediction Under the Ci PA Initiative. Clin. Pharmacol. Ther. 105, 466–475 (2019).

10. Mirams, G. R. et al. Simulation of multiple ion channel block provides improved early prediction of compounds’ clinical torsadogenic risk. Cardiovasc. Res. 91, 53–61 (2011).

11. Krogh-Madsen, T., Jacobson, A. F., Ortega, F. A. & Christini, D. J. Global Optimization of Ventricular Myocyte Model to Multi-Variable Objective Improves Predictions of Drug-Induced Torsades de Pointes. Front. Physiol. 8, 1–10 (2017).

12. Passini, E. et al. Drug□induced shortening of the electromechanical window is an effective biomarker for in silico prediction of clinical risk of arrhythmias. Br. J. Pharmacol. bph.14786 (2019).doi:10.1111/bph.14786

13. Lancaster, M. C. & Sobie, E. Improved Prediction of Drug-Induced Torsades de Pointes Through Simulations of Dynamics and Machine Learning Algorithms. Clin. Pharmacol. Ther. 100, 371–379 (2016).

14. Makkar, R. R., Fromm, B. S., Russell, T. S., Meissner, M. D. & Lehmann, M. H. Female Gender as a Risk Factor for Torsades de Pointes Associated With Cardiovascular Drugs. JAMA J. Am. Med. Assoc. 270, 2590 (1993).

15. Chorin, E. et al. Female gender as independent risk factor of torsades de pointes during acquired atrioventricular block. Hear. Rhythm 14, 90–95 (2017).

16. Flórez-Vargas, O. et al. Bias in the reporting of sex and age in biomedical research on mouse models. Elife 5, 1–14 (2016).

17. Vitale, C. et al. Under-representation of elderly and women in clinical trials. Int. J. Cardiol. 232, 216–221 (2017).

18. Ramirez, F. D. et al. Sex Bias Is Increasingly Prevalent in Preclinical Cardiovascular Research: Implications for Translational Medicine and Health Equity for Women. Circulation 135, 625–626 (2017).

19. Yang, P.-C., Kurokawa, J., Furukawa, T. & Clancy, C. E. Acute Effects of Sex Steroid Hormones on Susceptibility to Cardiac Arrhythmias: A Simulation Study. PLoS Comput. Biol. 6, e1000658 (2010).

20. Yang, P.-C. & Clancy, C. E. In silico Prediction of Sex-Based Differences in Human Susceptibility to Cardiac Ventricular Tachyarrhythmias. Front. Physiol. 3, 1–12 (2012).

21. O’Hara, T., Virág, L., Varró, A. & Rudy, Y. Simulation of the Undiseased Human Cardiac Ventricular Action Potential: Model Formulation and Experimental Validation. PLoS Comput. Biol. 7, e1002061 (2011).

22. Farrell, S. R., Ross, J. L. & Howlett, S. E. Sex differences in mechanisms of cardiac excitation-contraction coupling in rat ventricular myocytes. Am. J. Physiol. Circ. Physiol. 299, H36–H45 (2010).

23. Parks, R. J. & Howlett, S. E. Sex differences in mechanisms of cardiac excitation–contraction coupling. Pflügers Arch. -Eur. J. Physiol. 465, 747–763 (2013).

24. Parks, R. J., Ray, G., Bienvenu, L. A., Rose, R. A. & Howlett, S. E. Sex differences in SR Ca2+ release in murine ventricular myocytes are regulated by the cAMP/PKA pathway. J. Mol. Cell. Cardiol. 75, 162–173 (2014).

25. Colatsky, T. et al. The Comprehensive in Vitro Proarrhythmia Assay (CiPA) initiative — Update on progress. J. Pharmacol. Toxicol. Methods 81, 15–20 (2016).

26. Woosley, R., Heise, C., Gallo, T., Woosley, R. & Romero, K. QTDrug List. AZCERT at <https://crediblemeds.org/new-drug-list/>

27. Verkerk, A. O. et al. Gender Disparities in Cardiac Cellular Electrophysiology and Arrhythmia Susceptibility in Human Failing Ventricular Myocytes. Int. Heart J. 46, 1105–1118 (2005).

28. Fischer, T. H. et al. Sex-dependent alterations of Ca 2+ cycling in human cardiac hypertrophy and heart failure. Europace 18, 1440–1448 (2016).

29. Coppini, R. et al. Late Sodium Current Inhibition Reverses Electromechanical Dysfunction in Human Hypertrophic Cardiomyopathy. Circulation 127, 575–584 (2013).

30. Ferrantini, C. et al. Late sodium current inhibitors to treat exercise-induced obstruction in hypertrophic cardiomyopathy: an in vitro study in human myocardium. Br. J. Pharmacol. 175, 2635–2652 (2018).

31. Coppini, R. et al. Electrophysiological and Contractile Effects of Disopyramide in Patients With Obstructive Hypertrophic Cardiomyopathy. JACC Basic to Transl. Sci. 4, 795–813 (2019).

32. Furukawa, T. & Kurokawa, J. Regulation of cardiac ion channels via non-genomic action of sex steroid hormones: Implication for the gender difference in cardiac arrhythmias. Pharmacol. Ther. 115, 106–115 (2007).

33. Gaborit, N. et al. Gender-related differences in ion-channel and transporter subunit expression in non-diseased human hearts. J. Mol. Cell. Cardiol. 49, 639–646 (2010).

34. Rautaharju, P. M. et al. Sex differences in the evolution of the electrocardiographic QT interval with age. Can. J. Cardiol. 8, 690–5 (1992).

35. Yang, P.-C. et al. A Computational Pipeline to Predict Cardiotoxicity. Circ. Res. 126, 947–964 (2020).

36. Sims, C. et al. Sex, Age, and Regional Differences in L-Type Calcium Current Are Important Determinants of Arrhythmia Phenotype in Rabbit Hearts With Drug-Induced Long QT Type 2. Circ. Res. 102, 86–100 (2008).

37. Papp, R. et al. Genomic upregulation of cardiac Cav1.2α and NCX1 by estrogen in women. Biol. Sex Differ. 8, 5–8 (2017).

38. Tamari, I., Rabinowitz, B. & Neufeld, H. N. Torsade de pointes due to prenylamine controlled by lignocaine. Eur. Heart J. 3, 389–92 (1982).

39. Champeroux, P. et al. Prediction of the risk of Torsade de Pointes using the model of isolated canine Purkinje fibres. Br. J. Pharmacol. 144, 376–385 (2005).

40. Haverkamp, W. et al. Torsade de pointes induced by ajmaline. Z. Kardiol. 90, 586–590 (2001).

41. Vieweg, W. V. R. et al. Risperidone, QTc interval prolongation, and torsade de pointes: A systematic review of case reports. Psychopharmacology (Berl). 228, 515–524 (2013).

42. Harvey, P. A. & Leinwand, L. A. Oestrogen enhances cardiotoxicity induced by Sunitinib by regulation of drug transport and metabolism. Cardiovasc. Res. 107, 66–77 (2015).

43. Wenzel-Seifert, K., Wittmann, M. & Haen, E. QTc Prolongation by Psychotropic Drugs and the Risk of Torsade de Pointes. Dtsch. Aerzteblatt Online 108, 687–693 (2011).

44. Hasnain, M. et al. Quetiapine, QTc interval prolongation, and torsade de pointes: a review of case reports. Ther. Adv. Psychopharmacol. 4, 130–138 (2014).

45. Darpo, B. et al. Are women more susceptible than men to drug-induced QT prolongation? Concentration-QT c modelling in a phase 1 study with oral rac-sotalol. Br. J. Clin. Pharmacol. 77, 522–531 (2014).

46. Pokorney, S. D. et al. Dofetilide dose reductions and discontinuations in women compared with men. Hear. Rhythm 15, 478–484 (2018).

47. Thompson, K. A., Murray, J. J., Blair, I. A., Woosley, R. L. & Roden, D. M. Plasma concentrations of quinidine, its major metabolites, and dihydroquinidine in patients with torsades de pointes. Clin. Pharmacol. Ther. 43, 636–642 (1988).

48. Sauer, A. J. & Newton-Cheh, C. Clinical and Genetic Determinants of Torsade de Pointes Risk. Circulation 125, 1684–1694 (2012).

49. Huo, J., Wei, F., Cai, C., Lyn-Cook, B. & Pang, L. Sex-Related Differences in Drug-Induced QT Prolongation and Torsades de Pointes: A New Model System with Human iPSC-CMs. Toxicol. Sci. 167, 360–374 (2018).

