## Supplemental information for "Sex-specific classification of drug-induced Torsade de Pointes susceptibility using cardiac simulations and machine learning"

### *Corresponding author

### Detailed Methods

#### *1. Biophysical models and simulations*

We used the male and female human epicardial ventricular cardiomyocyte models developed by the Clancy lab[1] by incorporating experimentally determined sex- and hormone-specific differences in gene and protein expression into the O'Hara-Rudy model.[2] Some parameters of the baseline female model were modified to recapitulate observed functional sex differences in $Ca^{2+}$ handling.[3] Namely, we increased the maximal transport rate of the $Na^+$-$Ca^{2+}$ exchanger (NCX) by 15% in the female model and removed the originally introduced female-to-male differences in SERCA and $Na^+$/$K^+$-ATPase (NKA) formulations, in agreement with experimental measurements.[4,5]

To build the complete set of biomarkers used to train the ML classifiers, the virtual myocytes (with and without drug administration) were paced at a basic cycle length (BCL) of 500, 1,000, and 2,000 ms for 1,000 beats. Steady-state was confirmed as the intracellular $Na^+$ concentration had a beat-to-beat variation smaller than 1 µM. Each drug was simulated at multiple concentrations, ranging from 1 to 4 times their effective free therapeutic plasma concentration (EFTPC). The effects of each compound were simulated using a pore block model based on the available $IC_{50}$ values and Hill coefficients for various ion channels (a full list is available in **Table 1**). A total of 27 biomarkers (**Table 2**) were measured on the last simulated beat for each of the 12 conditions (4 concentrations x 3 pacing frequencies). Additional biomarkers included "direct" rather than simulated measures, such as $IC_{50}$ values for $I_{Kr}$, $I_{Na}$ (fast $Na^+$ current) and $I_{CaL}$ (L-type $Ca^{2+}$ current).

#### *2. Machine Learning*

A schematic diagram describing the complete workflow used in the generation of the ML classifiers for TdP risk is shown in **Fig 1**. First, the simulated recordings are inspected for the presence of early afterdepolarizations (EADs) or alternans in the last 3 beats. If proarrhythmic events are observed, e.g., EADs or repolarization failure that would alter the values of APD, the biomarkers for that specific drug-pacing frequency combination are imputed as follows: (1) the average value of each biomarker in the $TdP^+$ and $TdP^-$ groups is measured; (2) the drug receives the (maximum) minimum value of its TdP outcome group if the average value of its group is (larger) smaller than the value for the other group. For example, $TdP^+$ drugs have a larger average value for $APD_{90}$ than $TdP^-$ drugs. As a consequence, the $APD_{90}$ value of a $TdP^+$ drug producing EADs in a specific condition (e.g., Ibutilide at 4x EFTPC and 500 ms of BCL) will be the maximum $APD_{90}$ measured in that condition among all the $TdP^+$ drugs that do not produce EADs. To offset the differences of the sex-specific baseline models, the measured biomarkers in drug-free conditions are subtracted from those in response to drugs. Lastly, since biomarkers have different scales, each potential feature is standardized (i.e., the mean value is subtracted, and the difference is divided by the standard deviation) for better ML performance and interpretability.

In order to select the biomarkers contributing to the most accurate prediction of torsadogenic outcome, we applied a recursive feature elimination (RFE) algorithm. At each recursive step of the algorithm, a new classifier is trained with the available features and its regularization parameter is tuned to achieve the best performance. The predictive power of the model is quantified in the RFE algorithm using the Matthew's Correlation Coefficient (MCC) measured through Leave-One-Out Cross-Validation (LOO-CV). The features are then ranked based on their importance for the classification task, and the least important feature is discarded by the training dataset. The process is repeated until all the features are eliminated. The best candidate is the ML classifier characterized by the highest MCC using the smallest set of features. We calculated the area under the receiver operating characteristic curve (AUC) and the F1 score as additional performance metrics.

#### *3. Simulation and data analysis software, numerical method, and code availability*

The Yang and Clancy model[1] code was implemented in C++. We utilized the male model as is and modified the female model as described above. The ODEs were solved using a combination of Forward Euler and Rush-Larsen scheme, as done in the original O'Hara-Rudy model,[2] and implemented with a variable time step (dt = 0.025 or 0.005 ms). The data processing, RFE algorithm, and machine learning modeling were implemented in Python using the packages Numpy, Pandas, Scikit-Learn, and Hyperopt.

All simulations and data analyses were performed on a desktop server: HP Z2 Tower G4, Intel(R) Core(TM) i7-8700K @ 3.20GHz 6CPUs (12 threads) + 16GB; and a computing cluster with Intel(R) Xeon(R) CPU E5-2690 v4 @ 2.60GHz 28 CPUs (56 threads) + 132GB.

Source code and documentation are freely available at http://elegrandi.wixsite.com/grandilab/downloads and https://github.com/drgrandilab.

#### *4. Analysis of experimental $Ca^{2+}$ fluorescence data*

We retrospectively analyzed a dataset containing $Ca^{2+}$ transient properties in ventricular myocytes from hypertrophic cardiomyopathy patients (39 patients, 14 females and 25 males). Information on data collection can be found in the original published studies.[6–8] Statistical significance of differences was evaluated using an unpaired Student's *t*-test. Significance was set at an α-level of 0.05.

### Supplementary Tables

**Table S1**: Drug list with TdP risk category ($TdP^+$ in red, $TdP^-$ in green, intermediate risk in yellow), $IC_{50}$s and EFTPC. All concentrations are expressed in nM. A Hill coefficient (nH) of 1 was used unless a different nH is indicated in parenthesis.

| Drug | $I_{Kr}$ | $I_{NaL}$ | $I_{CaL}$ | $I_{Na}$ | $I_{to}$ | $I_{K1}$ | $I_{Ks}$ | EFTPC |
|---|---|---|---|---|---|---|---|---|
| Amiodarone 1 | 860 | | 1900 | 15900 | | | | 0.8 |
| Amiodarone 2 | 30 | | 270 | 4800 | | | | 0.5 |
| Bepridil 1 | 160 | | 1000 | 2300 | | | | 35 |
| Bepridil 2 | 33 | | 211 | 3700 | | | | 33 |
| Bepridil CiPA | 50 (0.9) | 1813.9 (1.4) | 2808.1 (0.6) | 2929.3 (1.2) | 8594 (3.5) | | 28628.3 (0.7) | 33 |
| Cilostazol | 13800 | | 91200 | 93700 | | | | 128 |
| Disopyramide | 14400 | | 1036700 | 168400 | | | | 742 |
| Dofetilide 1 | 30 | | 26700 | 162100 | | | | 2 |
| Dofetilide 2 | 5 | | 60000 | 300000 | | | | 2 |
| Dofetilide CiPA | 4.9 (0.9) | 753160.4 (0.3) | 260.3 (1.2) | 380.5 (0.9) | 18.8 (0.8) | 394.3 (0.8) | | 2 |
| Donepezil | 700 | | 34300 | 38500 | | | | 3 |
| Flecainide | 1500 | | 27100 | 6200 | | | | 753 |
| Halofantrine | 380 | | 1900 | 331200 | | | | 172 |
| Haloperidol 1 | 40 | | 1300 | 4300 | | | | 4 |
| Haloperidol 2 | 27 | | 1700 | 7000 | | | | 3.6 |
| Ibutilide | 18 | | 62500 | 42500 | | | | 140 |
| Methadone | 3500 | | 37400 | 31800 | | | | 507 |
| Moxifloxacin | 86200 | | 173000 | 1112000 | | | | 10960 |
| Procainamide | 272400 | | 389500 | 746600 | | | | 54180 |
| Quinidine 1 | 720 | | 6400 | 14600 | | | | 3237 |
| Quinidine 2 | 300 | | 15600 | 16600 | | | | 924 |
| Quinidine CiPA | 992 (0.8) | 9417 (1.3) | 51592.3 (0.6) | 12329 (1.5) | 3487.4 (1.3) | 39589919 (0.4) | 4898.9 (1.4) | 3237 |
| Sotalol | 111400 | | 193300 | 7013900 | | | | 14690 |
| Sotalol CiPA | 110600 (0.8) | | 7061527 (0.9) | 1140000000 (0.5) | 43143455 (0.7) | 3050260 (1.2) | 4221856 (1.2) | 14690 |
| Sparfloxacin | 22100 | | 88800 | 2555000 | | | | 1766 |
| Terodiline | 650 | | 4800 | 7400 | | | | 145 |
| Thioridazine 1 | 500 | | 3500 | 1400 | | | | 980 |
| Thioridazine 2 | 33 | | 1300 | 1830 | | | | 208 |
| Ajmaline | 1040 | | 71000 | 8200 | | | | 1500 |

| | | | | | | | | |
|---|---|---|---|---|---|---|---|---|
| Ceftriaxone | 445700 | | 153800 | 555900 | | | | 23170 |
| Cibenzoline | 22600 | | 30000 | 7800 | | | | 976 |
| Diazepam | 53200 | | 30500 | 306400 | | | | 29 |
| Diltiazem 1 | 13200 | | 760 | 22400 | | | | 122 |
| Diltiazem 2 | 17300 | | 450 | 9000 | | | | 122 |
| Diltiazem CiPA | 13150 (0.9) | 21868.5 (0.7) | 112.1 (0.7) | 110859 (0.7) | 2820000000 (0.2) | | | 122 |
| Duloxetine | 3800 | | 2800 | 5100 | | | | 16 |
| Lamivudine | 2054000 | | 54200 | 1571400 | | | | 19540 |
| Linezolid | 1147200 | | 105400 | 2644500 | | | | 59110 |
| Loratadine | 6100 | | 11400 | 28900 | | | | 0.4 |
| Mexiletine | 50000 | | 100000 | 43000 | | | | 4129 |
| Mexiletine CiPA | 28880 (0.9) | 8956.8 (1.4) | 38243.6 | | | | | 4129 |
| Mibefradil1 | 1700 | | 510 | 5600 | | | | 12 |
| Mibefradil2 | 1800 | | 156 | 980 | | | | 12 |
| Mitoxantrone | 539400 | | 22500 | 93500 | | | | 225 |
| Nifedipine1 | 44000 | | 12 | 88500 | | | | 8 |
| Nifedipine2 | 275000 | | 60 | 37000 | | | | 7.7 |
| Nitrendipine1 | 24600 | | 25 | 21600 | | | | 3 |
| Nitrendipine2 | 10000 | | 0.35 | 36000 | | | | 3.02 |
| Pentobarbital | 1433900 | | 299000 | 2686000 | | | | 5171 |
| Phenytoin1 | 147000 | | 21900 | 72400 | | | | 4360 |
| Phenytoin2 | 100000 | | 103000 | 49000 | | | | 4500 |
| Prenylamine | 65 | | 1240 | 2520 | | | | 17 |
| Propranolol | 2828 | | 18000 | 2100 | | | | 26 |
| Ribavirin | 967000 | | 622500 | 2997500 | | | | 27880 |
| Sitagliptin | 174700 | | 147100 | 1220800 | | | | 442 |
| Telbivudine | 422700 | | 713900 | 1095200 | | | | 19720 |
| Verapamil1 | 250 | | 200 | 32500 | | | | 88 |
| Verapamil2 | 143 | | 100 | 41500 | | | | 81 |
| Verapamil_CiPA | 288 | 7028 | 201.8 (1.1) | | 13429.2 (0.8) | 349000000 (0.3) | | 81 |
| Amitriptyline | 3280 | | 11600 | 20000 | | | | 41 |
| Astemizole | 4 | | 1100 | 3000 | | | | 0.3 |
| Chlorpromazine1 | 1500 | | 3400 | 3000 | | | | 38 |
| Chlorpromazine_CiPA | 929.2 (0.8) | 4559.6 (0.9) | 8191.9 (0.8) | 4535.6 (2) | 17616711 (0.4) | 9269.9 (0.7) | | 38 |
| Cisapride1 | 20 | | 11800 | 337000 | | | | 3 |
| Cisapride_CiPA | 10.1 (0.7) | | 9258076 (0.4) | | 219112.4 (0.2) | 29498 (0.5) | 81192862 (0.3) | 2.6 |
| Clozapine | 2300 | | 3600 | 15100 | | | | 71 |
| Dasatinib | 24500 | | 81100 | 76300 | | | | 41 |
| Desipramine | 1390 | | 1709 | 1520 | | | | 108 |
| Diphenhydramine | 5200 | | 228000 | 41000 | | | | 34 |

| | | | | | | | | |
|---|---|---|---|---|---|---|---|---|
| Droperidol | 60 | | 7600 | 22700 | | | | 16 |
| Fluvoxamine | 3100 | | 4900 | 39400 | | | | 377 |
| Imipramine | 3400 | | 8300 | 3600 | | | | 106 |
| Metronidazole | 1340200 | | 177900 | 2073200 | | | | 187000 |
| Nilotinib | 1000 | | 17500 | 13300 | | | | 172 |
| Ondansetron_CiPA | 1320 (0.9) | 19180.8 | 22551.4 (0.8) | 57666.4 | 1023378 | | 569807 (0.7) | 139 |
| Paliperidone | 780 | | 193900 | 109000 | | | | 69 |
| Paroxetine | 1900 | | 3900 | 9800 | | | | 14 |
| Pimozide1 | 40 | | 240 | 1100 | | | | 0.5 |
| Pimozide2 | 20 | | 162 | 54 | | | | 1 |
| Piperacillin | 3405100 | | 1226000 | 2433800 | | | | 1378000 |
| Propafenone | 440 | | 1800 | 1190 | | | | 241 |
| Quetiapine | 5800 | | 10400 | 16900 | | | | 33 |
| Raltegravir | 782800 | | 246700 | 824200 | | | | 7000 |
| Ranolazine_CiPA | 8270 (0.9) | 7884.5 (0.9) | | 68774 (1.4) | | | 36155020 (0.5) | 1948.2 |
| Risperidone1 | 260 | | 34200 | 43400 | | | | 2 |
| Risperidone2 | 150 | | 73000 | 102000 | | | | 1.81 |
| Saquinavir | 16900 | | 1900 | 12100 | | | | 130 |
| Sertindole1 | 33 | | 6300 | 6900 | | | | 2 |
| Sertindole2 | 14 | | 8900 | 2300 | | | | 1.59 |
| Solifenacin | 280 | | 4300 | 1500 | | | | 3 |
| Sunitinib | 1200 | | 33400 | 16500 | | | | 13 |
| Terfenadine1 | 50 | | 930 | 2000 | | | | 9 |
| Terfenadine2 | 8.9 | | 375 | 971 | | | | 9 |
| Terfenadine_CiPA | 23 (0.6) | 20056 (0.6) | 700.4 | 4803.2 | 239960.8 (0.3) | | 399754 (0.5) | 4 |
| Voriconazole | 490900 | | 414200 | 1550500 | | | | 7563 |

**Table S2:** List of biomarkers extracted from simulations and relative descriptions.

| Biomarker | Description |
|---|---|
| $APD_{90}$ | *Action potential duration at 90% repolarization* |
| $APD_{75}$ | *Action potential duration at 75% repolarization* |
| $APD_{50}$ | *Action potential duration at 50% repolarization* |
| $APD_{30}$ | *Action potential duration at 30% repolarization* |
| $V_{max}$ | *Peak voltage* |
| $V_{min}$ | *Diastolic voltage* |
| $dVdt_{max}$ | *Maximal upstroke velocity* |
| Plateau potential | *Average voltage between 10 and 50 ms after action potential initiation* |
| $I_{Na}$ max | *Peak of the sodium current* |
| CaT max | *Peak concentration of the calcium transient* |
| CaT min | *Diastolic intracellular calcium concentration* |
| $CaD_{80}$ | *Calcium transient duration at 80% return to baseline* |
| CaT tau | *Rate constant of decay of the calcium transient* |
| $Ca_i$ integral | *Integral of calcium transient* |
| $Na_i$ integral | *Integral of sodium transient ???* |
| $I_{Na}$ integral | *Integral of fast component of the sodium current* |
| $I_{NaL}$ integral | *Integral of late component of the sodium current* |
| $I_{to}$ integral | *Integral of transient outward potassium current* |
| $I_{CaL}$ integral | *Integral of L-type calcium current* |
| $I_{Kr}$ integral | *Integral of rapid delayed rectifier potassium current* |
| $I_{Ks}$ integral | *Integral of slow delayed rectifier potassium current* |
| $I_{K1}$ integral | *Integral of inward rectifier potassium current* |
| $I_{NaK}$ integral | *Integral of sodium/potassium-ATPase current* |
| $I_{pCa}$ integral | *Integral of calcium pump current* |
| $I_{NaCa}$ integral | *Integral of sodium/calcium exchanger current* |
| AP amplitude | *Action potential amplitude* |
| AP triangulation | *Difference between $APD_{80}$ and $APD_{30}$* |

### References


1. Yang, P.-C. & Clancy, C. E. In silico Prediction of Sex-Based Differences in Human Susceptibility to Cardiac Ventricular Tachyarrhythmias. *Front. Physiol.* **3**, 1–12 (2012).
2. O'Hara, T., Virág, L., Varró, A. & Rudy, Y. Simulation of the Undiseased Human Cardiac Ventricular Action Potential: Model Formulation and Experimental Validation. *PLoS Comput. Biol.* **7**, e1002061 (2011).
3. Parks, R. J., Ray, G., Bienvenu, L. A., Rose, R. A. & Howlett, S. E. Sex differences in SR Ca2+ release in murine ventricular myocytes are regulated by the cAMP/PKA pathway. *J. Mol. Cell. Cardiol.* **75**, 162–173 (2014).
4. Parks, R. J. & Howlett, S. E. Sex differences in mechanisms of cardiac excitation–contraction coupling. *Pflügers Arch. - Eur. J. Physiol.* **465**, 747–763 (2013).
5. Papp, R. *et al.* Genomic upregulation of cardiac Cav1.2α and NCX1 by estrogen in women. *Biol. Sex Differ.* **8**, 5–8 (2017).
6. Coppini, R. *et al.* Late Sodium Current Inhibition Reverses Electromechanical Dysfunction in Human Hypertrophic Cardiomyopathy. *Circulation* **127**, 575–584 (2013).
7. Ferrantini, C. *et al.* Late sodium current inhibitors to treat exercise-induced obstruction in hypertrophic cardiomyopathy: an in vitro study in human myocardium. *Br. J. Pharmacol.* **175**, 2635–2652 (2018).
8. Coppini, R. *et al.* Electrophysiological and Contractile Effects of Disopyramide in Patients With Obstructive Hypertrophic Cardiomyopathy. *JACC Basic to Transl. Sci.* **4**, 795–813 (2019).
